# Evidence of residual *Brugia malayi* infection in TAS-III cleared areas of coastal Odisha: Findings from Molecular Xenomonitoring

**DOI:** 10.64898/2026.07.17.739142

**Authors:** Kaushik Sanyal, Bhagyalaxmi Pradhan, Arundhuti Das, Dayeeta Bera, Arnav Pradhan, Udvas Ghorai, Shubhashisha Mohanty, Nilam Manoharrao Somalkar, Madhusmita Bal, P K Srivastava, Kalpana Baruah, Sanghamitra Pati

## Abstract

**Background:** Lymphatic filariasis (LF), caused by *Wuchereria (W) bancrofti*, *Brugia malayi* and *Brugia (B) timori*, remains a major public health problem known for its disfiguring and debilitating effects despite notable progress under the Global Program to Eliminate Lymphatic Filariasis (GPELF). Following recent national directives (2024) restructuring implementation units (IUs) to the block level, all TAS-cleared blocks now require re-evaluation using night blood surveys (NBS) or filarial test strip (FTS) kits. Molecular Xenomonitoring (MX) is recommended as a post-validation surveillance method for detecting active LF transmission in vectors. Therefore, an exploratory study was conducted in two coastal districts of Odisha, namely Jagatsinghpur and Puri, which cleared TAS-III, to investigate the presence of *B. malayi* and *W. bancrofti* infections in filaria-transmitting vectors.

**Methodology/Principal Findings:** Adult female mosquitoes were collected from three villages across Jagatsinghpur and Puri districts during March–April 2025. Species were identified morphologically, segregated and pooled. The pooled samples were screened for the presence of filarial DNA using species-specific PCR assays. Of 1,518 collected adult female mosquitoes, 22.6% were *Culex quinquefasciatus*, 18.3% were *Mansonia (Mn) annulifera*, and 4.3% were *Mansonia uniformis*. Among 33 pools screened, six pools were positive for *B. malayi* DNA (18.18% pool positivity; infection rate 2.88%, 95% CI: 1.23–5.74), in *Mn. annulifera*, whereas *W. bancrofti* was not detected.

**Conclusions/Significance:** The study provides the first molecular confirmation of *B. malayi* infection in *Mn. annulifera,* indicating a persistent Brugian transmission in Odisha even after the implementation of mass drug administration. The Study highlights MX as a sensitive, non-invasive surveillance tool for detecting residual infection in post-MDA and post-TAS settings. The study underscores the need for a standardized MX protocol and integration with entomological monitoring to sustain national LF elimination targets by 2027.

**Author Summary:** Lymphatic filariasis is a mosquito-borne disease that can cause severe, disfiguring swelling of the limbs and other lasting disabilities, and it remains a serious public health problem in India. National programmes in India are actively working to eliminate it by giving preventive medicines to communities, conducting surveys to confirm that transmission has ceased, and once an area passes, this treatment is withdrawn. However, it is not always clear whether the parasite has truly disappeared or is still lingering at very low levels. In this study, we searched for the parasite directly inside mosquitoes, an approach that does not require collecting blood from people. We sampled mosquitoes from two coastal districts of Odisha, India, that had already passed the required surveys and stopped treatment. We found the parasite *Brugia malayi* in *Mansonia* mosquitoes, the first time this has been shown using this molecular method in the region, although the more common filarial parasite was not detected. Our findings show that transmission can smoulder on even after an area is judged free of disease. Because India has no available rapid field test for this particular parasite, we argue that examining mosquitoes should become a routine safeguard for the country’s elimination effort.

## Introduction

Lymphatic filariasis (LF), is a neglected tropical vector-borne disease that causes immense suffering and socioeconomic deprivation among millions of people globally. The disease is caused by filarial nematodes such as *W. bancrofti*, *B. malayi* and *B. timori* transmitted through bites of infected mosquitoes. *B. timori* is restricted to Indonesia and not found in India. LF leads to significant chronic morbidity and disability, such as lymphedema, hydrocele, affecting large population in low and middle-income countries [1–3]. At present, more than half of the global LF burden is concentrated in Southeast Asia, with India alone contributing over 23 million cases and placing 650 million people at risk [4,5]. India accounts ∼62% (404.3 million people) at risk group and reports approximately 619, 000 cases of lymphedema and 150, 708 hydrocele cases [6,7]. Notably, four states, such as Odisha, Uttar Pradesh, Telangana and Bihar, collectively contribute 60% of lymphedema cases and 80% of hydrocele cases [7–9].

Recognising the global importance of eliminating LF and in pursuit to fulfil the commitment of World Health Assembly (WHA) resolution of 1997, the World Health Organization (WHO) launched the GPELF in 2000 with dual strategies of Mass Drug Administration (MDA) using a combination of either Ivermectin or Diethylcarbamazine with Albendazole and Morbidity Management Disability Prevention (MMDP) [10,11]. This integrated approach of the GPELF program has yielded substantial progress, contributing 74% global reduction in LF prevalence, while complete elimination remains an unfinished goal. Initially WHO target was to eliminate LF by 2020, which was subsequently extended to 2030, aligning with Sustainable Development Goal (SDG), while India has set an ambitious national goal to achieve elimination by 2027 [7]. Historically, MDA for LF towards elimination strategy was initiated in India as pilot during 1996-1997 across 13 districts in 7 states, using a single-dose Diethylcarbamazine (DEC) regimen [12]. The pilot project was expanded in 2002 to 31 districts, and selected areas adopted the DEC-Albendazole (ALB) combination therapy [2,12]. A nation-wide implementation began in 2004 with the goal of elimination through the Accelerated Plan for Elimination of LF (APELF), continuing till 2024 [7]. To further accelerate progress, India introduced triple drug therapy with Ivermectin + DEC + ALB (IDA) in 2018 [13,14].

The outcome of MDA has relied on parasitological and immunological assessments, including microfilaria detection in both the vector species and circulating filarial antigenemia (CFA) detection for *W. bancrofti*, and antibody detection for *B. malayi* [9]. Also, for post MDA surveillance, Transmission Assessment surveys (TAS) serve as a standardized evaluation method, employing Immuno-chromatographic tests (ICT) card-like rapid diagnostic tests [6,11,15–18].

Currently, out of 345 endemic districts [7], 170 continue MDA implementation, 138 cleared TASI, discontinued MDA, and 37 are under evaluation. The minimum effective MDA coverage required for the interruption of transmission is >90% against the eligible population, now set by the National Program [7]. Notably, the coverage has increased from 72.42% in 2004 to 87.25 % in 2019 [19–26]. Despite these achievements, sustained success at the national level will require a systematic mapping of LF vectors at the district levels in post-MDA and post-TAS settings to detect active transmission [7].

Mosquito dissection remains one of the traditional gold-standard techniques for estimating infection rates and parasite densities. However, performing the large number of dissections required for reliable estimates is costly, time-consuming, and labour-intensive, particularly in settings with low infection prevalence [27]. Molecular xenomonitoring (MX), a sensitive PCR-based approach, detects parasite DNA in mosquitoes and serves as a proxy indicator of ongoing lymphatic filariasis (LF) transmission and recent human infection [11]. MX is recognized by WHO as one of the recommended surveillance strategies and can detect microfilariae in pooled mosquito samples of 50–100 specimens [28–30]. It complements Transmission Assessment Surveys (TAS) by confirming LF infection when prevalence becomes undetectable by conventional methods and supports decisions regarding the premature cessation of mass drug administration (MDA) [29]. According to WHO, the standard Vector infection thresholds to decide MDA cessation are < 0.25% for *Culex,* < 1% for *Anopheles*, <0.5% for *Mansonia* and <0.1% for *Aedes* species [11]. Nevertheless, confirmation of ongoing transmission remains challenging due to the lack of standardized sampling strategies and the need for further research to establish definitive DNA-prevalence thresholds in vector populations [29–31]

MX studies have been undertaken for bancroftian infection in different Indian states viz., Tamil Nadu, Kerala, Karnataka, Madhya Pradesh, Puducherry and Odisha [29,32–37]. It has been recommended as a proxy tool for monitoring in both pre and post-MDA settings with diverse objectives [32–37]. In a post-MDA and post-TAS setting, the primary goal of surveillance is to identify a potential source of recrudescence through targeted sampling with strong entomomolecular diagnostic capacity. However, the implementation of MX in operational settings needs an evaluation of its cost-effectiveness and feasibility to TAS, considering the availability of suitable laboratory infrastructure and trained personnel [27,29,38–40].

*W. bancrofti* is main LF infection in Odisha, transmitted by vector *Culex quinquefasciatus* across 20 districts; however, *B. malayi* cases have been reported in coastal districts like Bhadrak and Baleswar, where presence of perennial ponds with abundant aquatic vegetation favours *Mansonia* breeding due to larval attachment to plant roots [37,41]

With multiple rounds of MDA, the overall microfilaria (mf) rate has reduced to 0.68% by 2015, though the actual drug compliance has been lower than the reported coverage during MDA, which needs to be bridged for programme’s success [5,42]. Based on reduced mf rate and more than 5 rounds of MDA with compliance over 65%, TAS with FTS is done and if the district clears, MDA is stopped and the district (Implementation unit) undergoes post MDA surveillance.

Based on the facts of presence of both the infection of LF prevailing in Odisha, this exploratory study aimed to gain insights and map the spatial distribution of LF transmitting vectors to identify the potential foci of ongoing transmission across the TAS-cleared districts using MX survey [43]. The study sought to generate data about the vector abundance, diversity and entomological risk map to guide an evidence-based decision-making for LF elimination. Additionally, this study also aimed to evaluate field-based vector sampling strategies for future longitudinal studies in Odisha.

## Materials and Method

### Study site

The study was conducted in three villages of two districts-Jagatsinghpur and Puri. Two villages were Bhimanasi (Latitude 20.308401° N, Longitude 86.639152° E) and Musadia (Latitude 20.321404° N, Longitude 86.653055° E) selected in Jagatsinghpur and one village, Patasundarpur (Latitude 20.039341° N, Longitude 86.196039° E), from Puri. The study sites were located in the coastal part of Odisha[44]. Both districts have rice growing alluvial plains, surrounded by river basin. Persistent breeding habitats were present in all three villages, including ponds rich in aquatic vegetation, ditches with standing water, damp and swampy low-lying areas and clogged drains with stagnant water[45]. These conditions supported the breeding of both *Cx. quinquefasciatus* and *Mansonia* species. Unlined drains, standing water sources, and high human-vector contact because of poor housing infrastructure were common risk factors for LF transmission in the villages’ ecological setting as reported earlier also[46]. In both these selected districts, MDA was started in 2004, and TAS was cleared in 2017 (unpublished data).

### Mosquito sample collection

The study was conducted during hot season between March and April 2025. The study sites were selected based on the presence of breeding potential and epidemiological record. Oral consent was obtained from households with reported LF cases. Adult female mosquitoes were collected using oral aspirator[6] from the three selected sites during the night hours (22:00 - 24:00 hours) from randomly selected households, cattlesheds nearer to breeding habitats (swampy and riverine areas). A total of 1518 female adult mosquitoes were collected from all sites and brought to laboratory for further molecular analysis.

### Morphological identification

Adult mosquitoes were morphologically identified under Stereozoom microscope (Metzer-M) using standard taxonomic keys of Tyagi et al [47].The species wise segregation of adult female mosquitoes was done. The mosquitoes were pooled (20-25 specimens per pool) keeping each species, household, name of village and collection date separate and stored at -20°C before molecular analysis at ICMR-RMRC Bhubaneswar laboratory. Some pools had <20 mosquitoes due to limited availability of species.

### DNA extraction and PCR analyses

DNA was extracted from each mosquito pool using Qiagen Dneasy Blood and Tissue Kit (catalogue no. 69506; Qiagen, Hilden, Germany) in accordance with manufacturer’s instructions. Conventional PCR was conducted (20 µl reaction volume) to detect the infection of *W. bancrofti* using a primer pair targeting SspI repeat region, forward primer-5’- CAAAGTAGCGTAAGGGAATTG-3’, reverse primer-5′- CCCTCACTTACCATAAGACAAC-3 and for *B. malayi Hha*I repeat region is targeted, forward primer-5’-CTTCATTAGACAAGGATATTGGTTC-3’, reverse primer-5’- GACAACACAATACACGACCAG-3^’^ [48]. Amplification was carried out with activation stage at 95°C for 5 min, followed by a 30-cycle amplification of 30s at 95°C, then at 56°C for 30s and at 70°C for 5 min, with the final extension step at 70°C for 5 min. The final PCR product was visualized on 2% agarose gel and the samples were considered positive for parasite infection when species-specific bands with expected size were observed.

### Determination of pool positivity and maximum likelihood estimation

Data from female adult mosquito collection and PCR analysis were used to characterize major LF vector abundance and proportion of pool positivity by species. The pool positivity was determined by dividing the total number of positive tested pools for filarial infection by the total number of pools screened. Furthermore, infection rates (IR) were calculated using Maximum Likelihood Estimation (MLE) [32] a statistical approach that estimates the most likely prevalence of infection based on the number of pools tested, the number of mosquitoes in each pool, and the number of positive pools. Additionally, we created a graph showing infection rate by different mosquito species, including confidence intervals.

## Results

### Distribution of Mosquitoes and their abundance

During the study, aspirator-based sampling was deployed near mixed dwellings of households (HH) and cattle sheds in each village, of total 1518 collected adult female mosquitoes, of which 344 (22.6%) were *Cx. quinquefasciatus*, 279 (18.3%) were *Mn. annulifera* and 65 (4.28%) *Mn. uniformis* (Table 1, Fig 1). Although having a smaller sample size among all the studied sites, we found *Cx. vishnui*, *Cx. quinquefasciatus* and *Mn. annulifera* were predominant in both villages of Jagatsinghpur district while *Cx. tritaeniorhynchus*, *Mn. annulifera* and *Cx. gelidus* were found to be more in Patasundarpur village in Puri (Table 1). *Mn. annulifera* were detected across all sites with Musadia exhibiting the highest concentration. These findings indicated vector preferences are location specific.

**Table 1.**
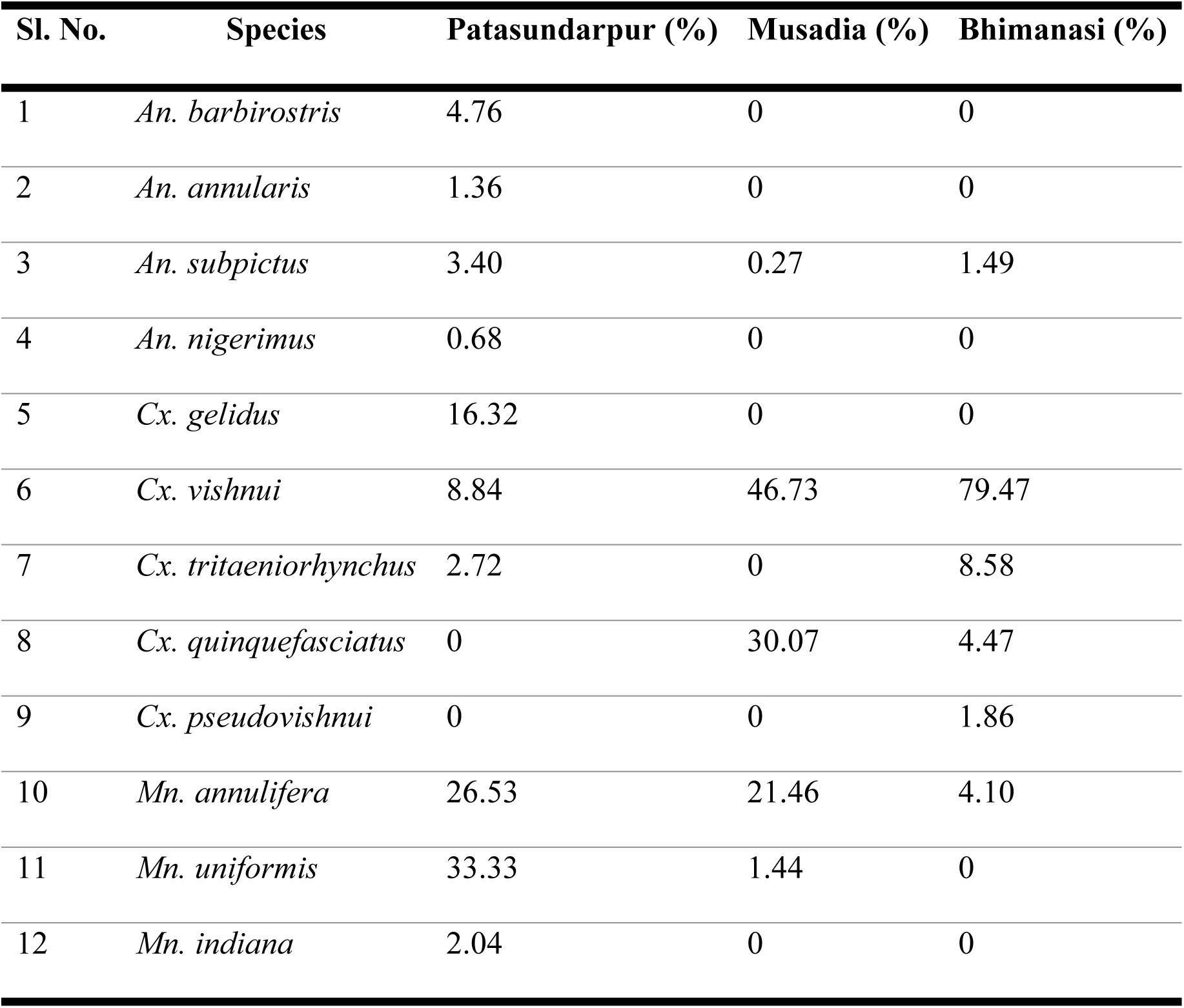
Species-wise distribution of adult mosquitoes collected across villages in Puri and Jagatsinghpur districts.

| Sl. No. | Species | Patasundarpur (%) | Musadia (%) | Bhimanasi (%) |
| --- | --- | --- | --- | --- |
| 1 | *An. barbirostris* | 4.76 | 0 | 0 |
| 2 | *An. annularis* | 1.36 | 0 | 0 |
| 3 | *An. subpictus* | 3.40 | 0.27 | 1.49 |
| 4 | *An. nigerimus* | 0.68 | 0 | 0 |
| 5 | *Cx. gelidus* | 16.32 | 0 | 0 |
| 6 | *Cx. vishnui* | 8.84 | 46.73 | 79.47 |
| 7 | *Cx. tritaeniorhynchus* | 2.72 | 0 | 8.58 |
| 8 | *Cx. quinquefasciatus* | 0 | 30.07 | 4.47 |
| 9 | *Cx. pseudovishnui* | 0 | 0 | 1.86 |
| 10 | *Mn. annulifera* | 26.53 | 21.46 | 4.10 |
| 11 | *Mn. uniformis* | 33.33 | 1.44 | 0 |
| 12 | *Mn. indiana* | 2.04 | 0 | 0 |

**Fig 1.**
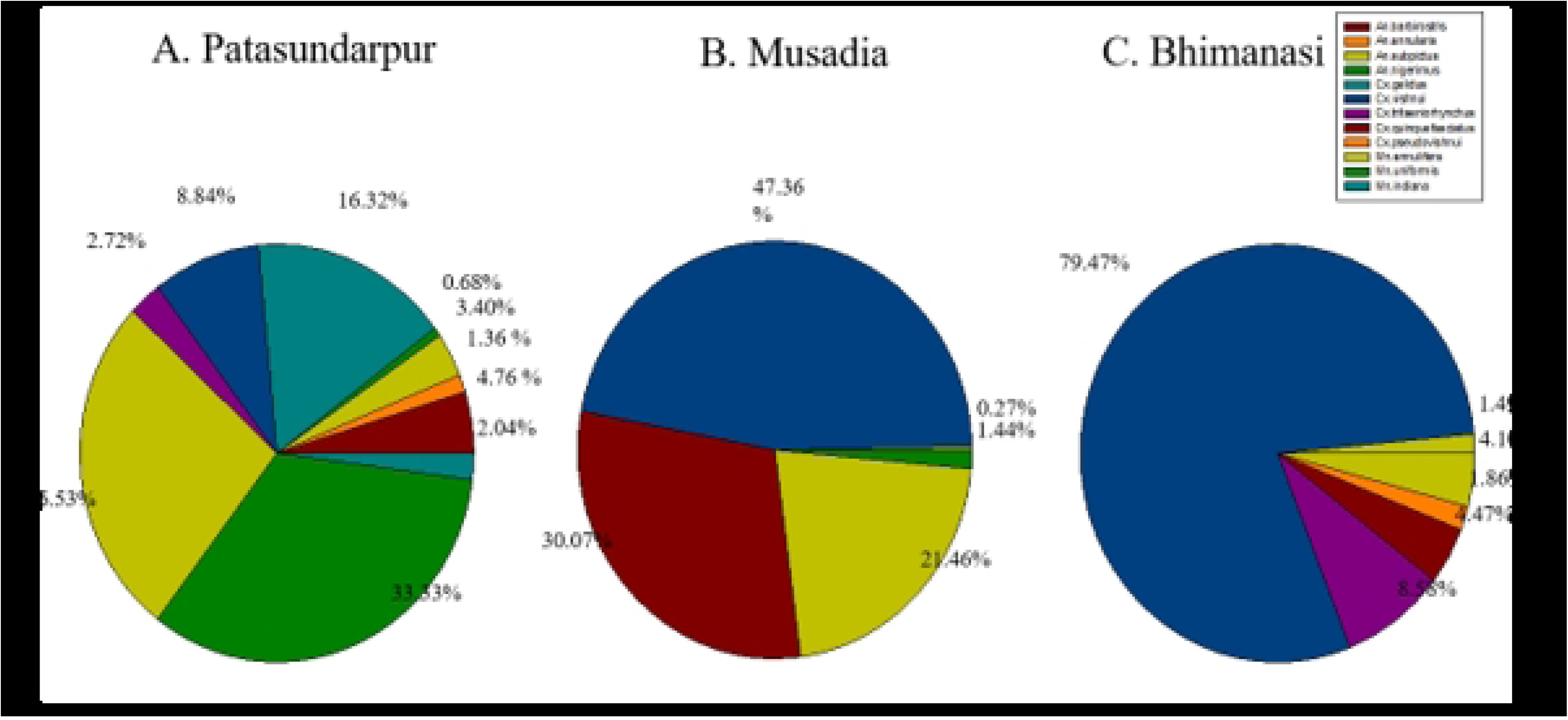
Species composition of mosquito vectors collected from three villages (Patasundarpur, Musadia, and Bhimanasi), showing relative abundance of different *Anopheles, Culex*, and *Mansonia* species.

### Detection of *B.malayi* in PCR assay and LF positive Mosquitoes

A total of 33 pools were analysed using conventional PCR. All pools were tested negative for *W. bancrofti* infection while 6 pools were detected positive for *B.malayi* DNA as confirmed by 2% agarose gel electrophoresis (Fig 2)*. The six B.malayi* pools represent a pool positivity 18.18% with an infection rate 2.88%, confidence intervals 95%(CI): 1.23-5.74 exceeding provisional WHO threshold [34] (Fig 3). All *Brugia* positives detections were from *Mn. annulifera* showed a species-specific positivity rate was 46.15% (6/13).

**Fig 2.**
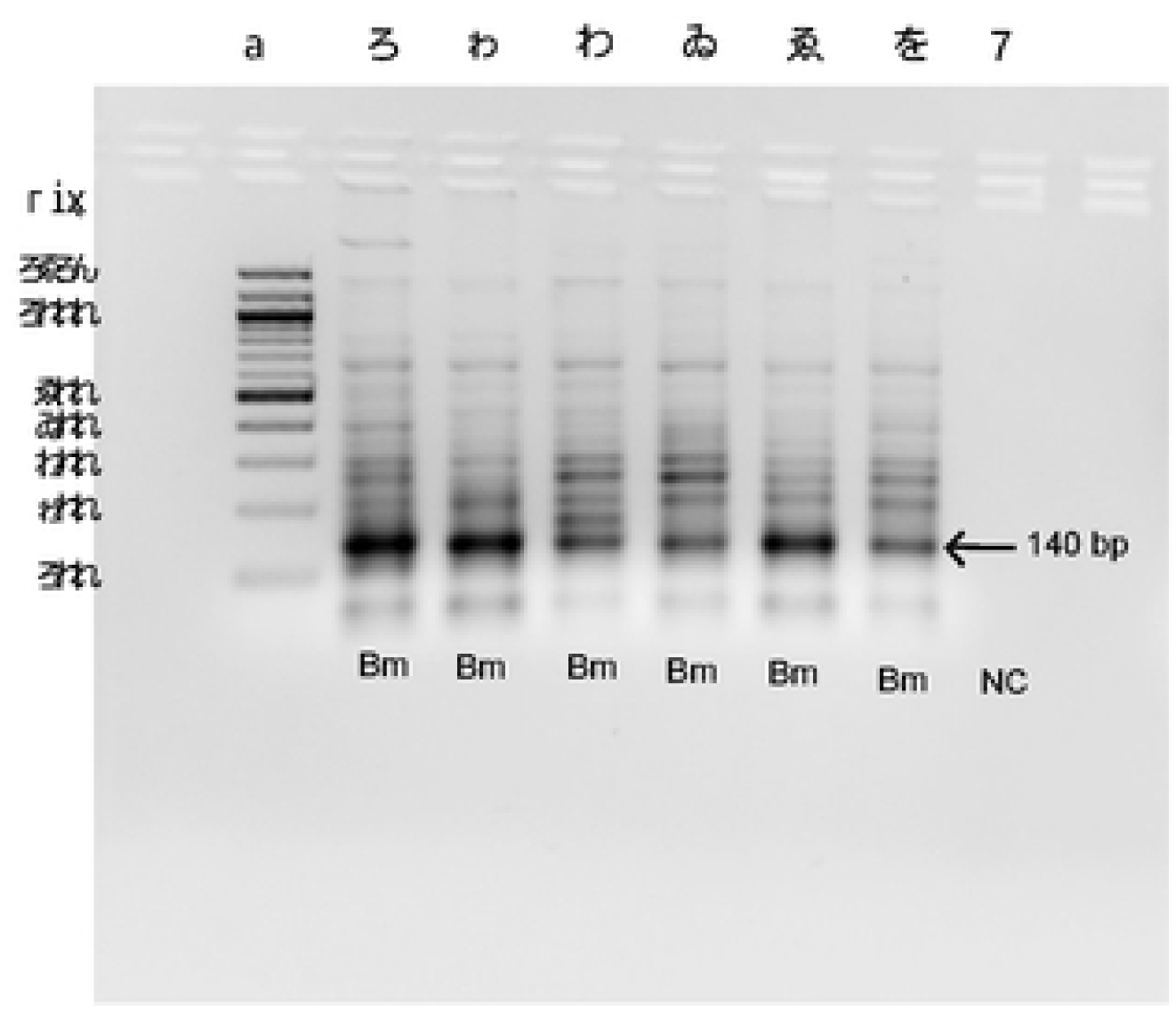
Optimized PCR assay for detection of *B. malayi* DNA^34^ in 2% agarose gel electrophoresis showed the amplified Hha1 gene of *B. malayi.* Lane M: 1.5 Kb DNA ladder; Lanes 1-6 represent positive *B.malayi* DNA samples, Lane 7 represents negative control.

**Fig 3.**
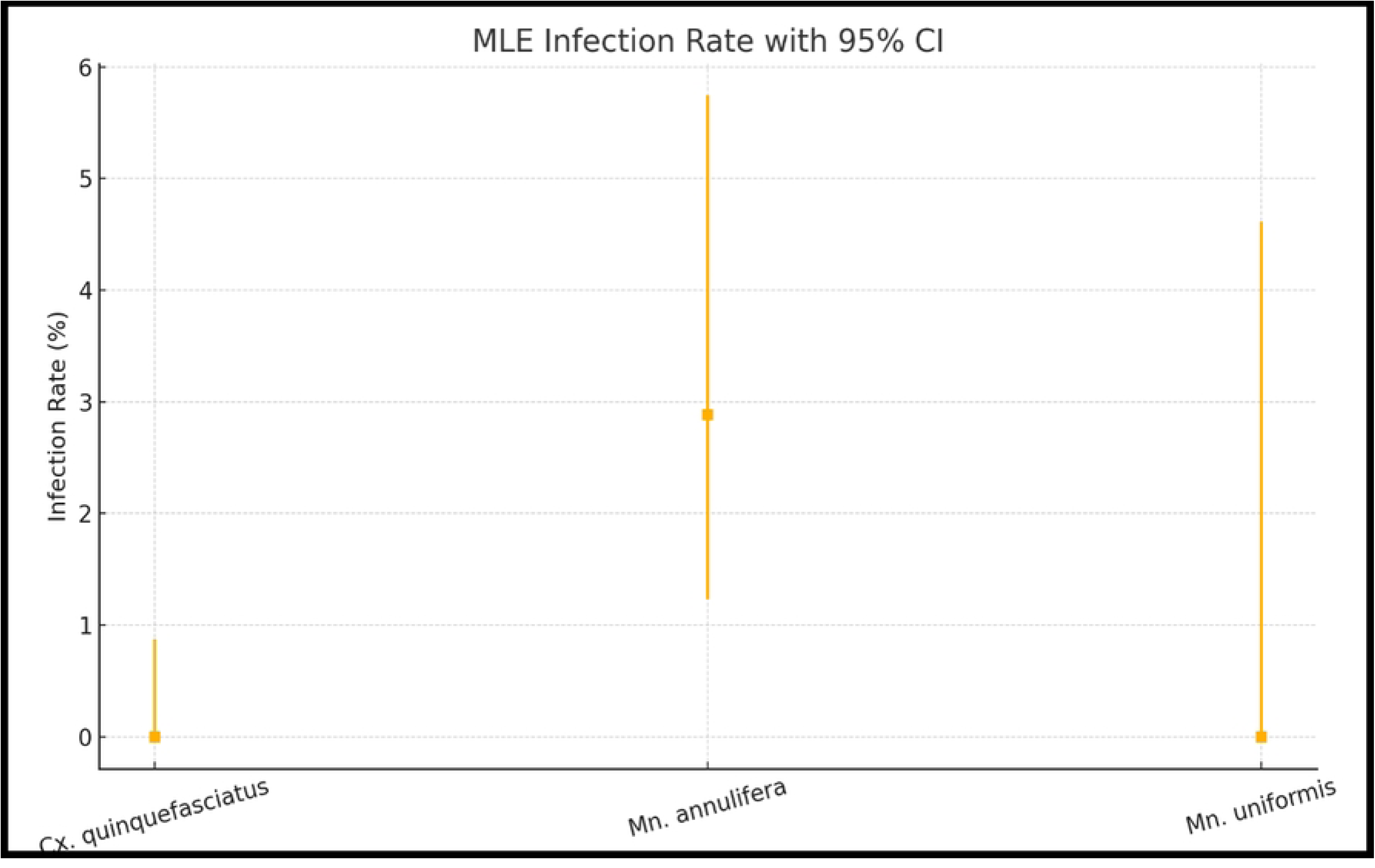
Maximum likelihood estimates (MLE) of infection rates in mosquito species (*C*. *quinquefasciatus*, *Mn*. *annulifera*, and *Mn*. *uniformis*) collected during the pilot molecular xenomonitoring study

### Maximum Likelihood Estimation (MLE) Rate in Mosquitoes

The MLE infection rate of mosquito sample revealed that *Mn. annulifera* exhibited 2.88% of mean infection rate of *B. malayi* with a wide 95% CI (∼1% to ∼6%) suggesting moderate level of infection presence but some uncertainty likely due to possibly limited sample size. In contrast, *Mn. uniformis* showed low estimated infection (∼0–1%) but with a broad CI (∼0– 4.5%), suggesting low infection but less precision in the estimate (Fig 3). The persistence of filarial transmission at low levels, highlights the importance of both vector and poor/non-vector species for ongoing transmission necessitating the vector surveillance for proper vector control programme.

## Discussion

This study represents the first exploratory report documenting *Brugia malayi* infection in *Mansonia* species from Odisha confirmed through molecular xenomonitoring. The investigation evaluated the abundance and spatial distribution of predominant vector species, their incrimination status, and the pattern of ongoing LF transmission using a molecular xenomonitoring (MX) approach. The results indicated that the nocturnal *Culex spp.* were the most abundant in the surveyed areas; however, none of its pools were detected positive for *W.bancrofti* infection. In contrast, *Mansonia* species exhibited pool positivity for *B. malayi*, differing from earlier observation in Balasore district [37]. The overall pool positivity rate in our study (18.18%) for *B.malayi* suggests a substantial nematode infection in mosquitoes and indicates a continuing risk of transmission in the study area as was also described in a study of Madhya Pradesh [36].

The absence of parasite DNA detection in *Culex* pools should be interpreted cautiously, as they may reflect limited sampling intensity rather than the true absence of infection. The predominantly nocturnal, exophilic biting behaviour of *Culex* spp. with modest peaks during dusk, midnight and pre-dawn hours may partly explain the low detection of *W. bancrofti*. The vector collections from structures with low density or in non-peak biting hours may miss infected mosquitoes, resulting in false negative findings. Therefore, expanding mosquito sampling effort both spatially and temporally along with molecular screening of larger sample sizes, is necessary for reliable detection of low prevalence infections [34,36]. Additionally, understanding ecological determinants is essential for identifying vector breeding preferences and ensuring the sustainability of a long term control efforts [10]

Previous studies across India have consistently reported high pool positivity rates for *W. bancrofti* in *Culex quinquefasciatus* during post MDA period [29,32–37] suggesting it as a complementary tool to ensure absence of transmission. Similar, investigations conducted in Alappuzha, Kerala, an endemic hotspot for both parasites, have reported the re-emergence of *Brugian* infection despite successful vector control efforts [34]. In addition, a recent study in Yadgir district of Karnataka demonstrated *W.bancrofti* infection in both vector (*Culex quinquefasciatus*)and non-vector (*An subpictus* and *Culex gelidus*)species [35]. Studies have revealed that Odisha is also a focus for both *Brugian and Bancroftian* filariasis [32,33,35,47]. The detection of multiple mosquito species from three clusters, including known vector and non-vector species, underscores the complexity of vector ecology, ongoing parasite circulation in the local environment, however, it does not imply active transmission by non-vector species like *Mansonia* sp. These findings support the need for strengthened integrated entomological surveillance, targeted vector control, and enhanced community awareness [36]. Periodic implementation, review, and validation of MX surveillance are therefore recommended to ensure accurate monitoring of LF transmission dynamics.

MX is a highly sensitive measure of parasite presence in vector populations during and after MDA application even after the occurrence of infection is <1% [10,16,49–53]. In the present study, MX was able to detect the presence of parasite DNA, though it could not detect the infective L3 stage. The detection of *B. malayi* in *Mansonia* spp should therefore be interpreted as evidence of local parasite circulation rather than direct confirmation of infective larvae. Confirmation of infective stages would require RT-PCR assays targeting L3 stage-specific gene expression [29]. Although multicentric and stage specific RT-PCR assay was developed for detecting infective *W. bancrofti* (L3) larvae stage in *C. quinquefasciatus* [53], operational challenges remain, including sample processing cold-chain requirements, RNA instability, and field-to-laboratory transport constraints. Given the detection of *Brugian* DNA in our present study, there is a critical need to develop and deploy a *Brugian* infective stage (L3) specific quantitative RT-PCR method, as demonstrated in limited global studies [54].

Despite TAS III clearance, the observed pool positivity for *Brugia* among *Mansonia* species suggests the possibility of continued LF transmission in post-MDA period, aligning with earlier findings from TAS-III cleared evaluation unit in Odisha [32]. Comparable patterns have been reported in Sri Lanka where vector infection (5.7%) and CFA prevalence (3%) were observed two years after WHO validation of LF elimination [50] emphasizing the need for vector mapping in TAS-cleared regions. Studies from Tanzania’s Mafia Islands and American Samoa have similarly shown that MX detected ongoing transmission risk even when TAS suggested interruption [52]. Although Puri and Jagatsinghpur districts achieved mf rates below 1% and cleared TAS in 2017 (unpublished data), the present post-TAS MX survey detected a *B. malayi* infection rate of 2.88% (95% CI above the WHO threshold), indicating the need for localized surveys or targeted treatment rounds [55].

According to WHO, detection of *W. bancrofti* relies on both Mf detection and antigen (Ag)-based assay, while *B. malayi* surveillance due to the lack of *Brugia*-specific antigen detection kit in India depends primarily on Mf detection which are labour-intensive and constrained by parasite periodicity [56] highlighting the need for a reliable *Brugia-*specific point-of-care diagnostic tool for remapping in endemic regions prior to elimination. National guidelines also recommend continuing post-elimination surveillance for atleast 10 years using MX in alongside standard epidemiological surveys to verify post-validation status and detect any resurgence or potential of transmission [15]. In Indian LF elimination programme implementation units (IU) have narrowed down from district-level to block-level, covering approximately 1.5 lakh population [15]. Further, the monitoring framework has also been streamlined from four sentinel and four random sites per district to one sentinel and two random sites increasing the importance of strengthened vector surveillance for smaller unit areas. The revised guidelines have changed the sample sizes and target groups: NBS samples have been reduced from 500 to 300 per site; TAS under DA regimen continue to target children aged 6– 7 years, while under IDA, adults above 20 years are included with larger sample sizes (∼3150) and mf testing among FTS-positive individuals [7] Despite these refinements, standardized MX sampling protocol is needed in national or global guidance. WHO has lowered the antigenemia threshold for TAS clearance from <2% to <1%, and the same need for aligned under national guidelines [7] (Fig 4).

**Fig 4.**
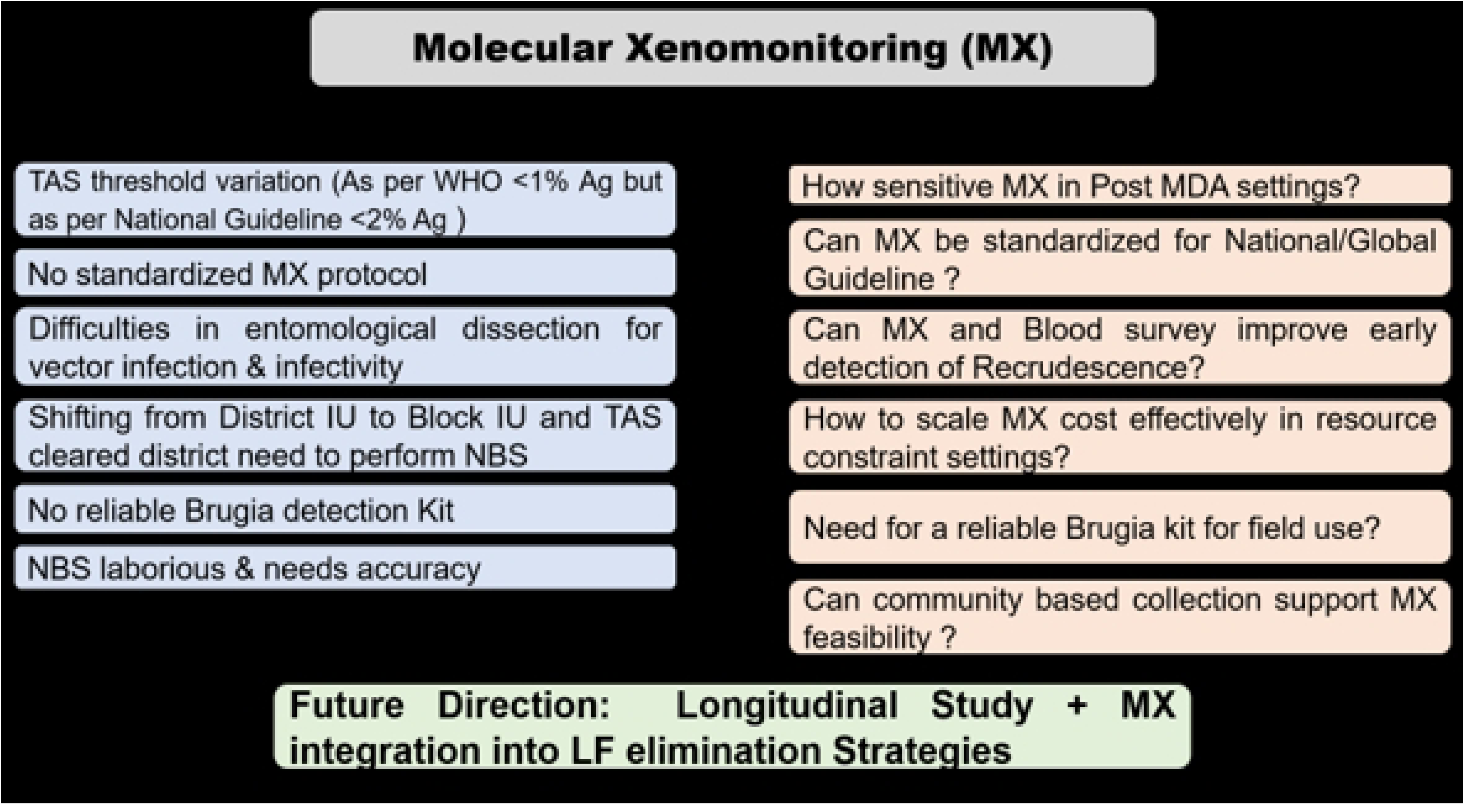
Challenges and research priorities for integrating MX into LF elimination, highlighting gaps, with future directions focused on longitudinal studies and community-based approaches.

Despite limitations of small sample size, non-standardized sampling and absence of infective L-3 larval stage confirmation, this study indicates possible ongoing Brugian transmission after TAS clearance. Periodic MX surveillance, integrated with epidemiological survey and strengthened vector monitoring will be important to prevent resurgence and sustain LF elimination even after achieving it as also recommended by WHO guidelines 2025 [11].

## Conclusion

This study provides the first evidence of *B. malayi* infection in *Mansonia* species in Odisha, indicating possible ongoing LF transmission despite TAS clearance. Because TAS currently relies on *W. bancrofti*–specific FTS kits and no diagnostic kit is available in India for *B. malayi*, MX emerges as a sensitive and operationally valuable complementary tool for detecting residual infection and interpreting post-MDA transmission dynamics. MX also contributes important insights into vector abundance and ecology, supporting targeted vector management. Despite sampling and stage-confirmation limitations, the findings highlight the need for expanded longitudinal MX surveillance. Institutionalizing MX within the national programme will be essential for early detection, targeted response, and prevention of LF resurgence beyond 2027.

## Acknowledgements

The authors sincerely thank the Director of Public Health, Government of Odisha, the State Vector Borne Disease Control Programme (NVBDCP), and the district health authorities of Jagatsingpur and Puri for their support in facilitating the field activities. We also express our heartfelt gratitude to the community members and local leaders of the study villages for their collaboration and participation throughout the study.

## Author’s contributions

**Conceptualization**: Kaushik Sanyal, Shubhashisha Mohanty, Nilam Manoharrao Somalkar, Madhusmita Bal, Kalpana Baruah, Sanghamitra Pati

**Data curation**: Kaushik Sanyal, Bhagyalaxmi Pradhan, Udvas Ghorai

**Formal analysis**: Kaushik Sanyal, Bhagyalaxmi Pradhan, Arundhuti Das

**Methodology**: Kaushik Sanyal, P K Srivastava, Kalpana Baruah

**Investigation**: Kaushik Sanyal, Bhagyalaxmi Pradhan

**Project administration**: Kaushik Sanyal

**Resources**: Kaushik Sanyal, Sanghamitra Pati

**Supervision**: Kaushik Sanyal

**Validation**: Kaushik Sanyal, Arundhuti Das, Shubhashisha Mohanty, Nilam Manoharrao Somalkar, P K Srivastava, Kalpana Baruah

**Visualization**: Kaushik Sanyal, Bhagyalaxmi Pradhan, Dayeeta Bera, Arnav Pradhan

**Writing – original draft**: Kaushik Sanyal, Bhagyalaxmi Pradhan, Arundhuti Das

**Writing – review & editing**: Kaushik Sanyal, Arundhuti Das, Shubhashisha Mohanty, Madhusmita Bal, P K Srivastava, Kalpana Baruah, Sanghamitra Pati

## Financial Disclosure Statement

This work was supported by an intramural research grant from the Indian Council of Medical Research (Project Sanction Order No. INTR-IM-2024-00120), awarded to Kaushik Sanyal. The funder had no role in study design, data collection and analysis, decision to publish, or preparation of the manuscript.

## Competing Interests

The authors declare no competing interests.

## Notes

### Competing Interest Statement

The authors have declared no competing interest.

